# An engineered receptor-binding domain improves the immunogenicity of multivalent SARS-CoV-2 vaccines

**DOI:** 10.1101/2020.11.18.388934

**Authors:** Brian D. Quinlan, Wenhui He, Huihui Mou, Lizhou Zhang, Yan Guo, Jing Chang, Shoujiao Peng, Amrita Ojha, Rubens Tavora, Mark S. Parcells, Guangxiang Luo, Wenhui Li, Guocai Zhong, Hyeryun Choe, Michael Farzan

**Affiliations:** Department of Immunology and Microbiology, The Scripps Research Institute, Jupiter, FL 33458, USA; Department of Animal and Food Sciences, University of Delaware, Newark, DE 19716, USA; Department of Microbiology, University of Alabama at Birmingham School Of Medicine, Birmingham, AL 35294, USA; National Institute of Biological Sciences, Tsinghua Institute of Multidisciplinary Biomedical Research, Tsinghua University, Beijing, China; Scripps Research | SZBL Chemical Biology Institute, Shenzhen Bay Laboratory (SZBL), Shenzhen, China; School of Chemical Biology and Biotechnology, Peking University Shenzhen Graduate School, Shenzhen, China

## Abstract

The SARS-coronavirus 2 (SARS-CoV-2) spike (S) protein mediates viral entry into cells expressing the angiotensin-converting enzyme 2 (ACE2). The S protein engages ACE2 through its receptor-binding domain (RBD), an independently folded 197-amino acid fragment of the 1273-amino acid S-protein protomer. The RBD is the primary SARS-CoV-2 neutralizing epitope and a critical target of any SARS-CoV-2 vaccine. Here we show that this RBD conjugated to each of two carrier proteins elicited more potent neutralizing responses in immunized rodents than did a similarly conjugated proline-stabilized S-protein ectodomain. Nonetheless, the native RBD expresses inefficiently, limiting its usefulness as a vaccine antigen. However, we show that an RBD engineered with four novel glycosylation sites (gRBD) expresses markedly more efficiently, and generates a more potent neutralizing responses as a DNA vaccine antigen, than the wild-type RBD or the full-length S protein, especially when fused to multivalent carriers such as an *H. pylori* ferritin 24-mer. Further, gRBD is more immunogenic than the wild-type RBD when administered as a subunit protein vaccine. Our data suggest that multivalent gRBD antigens can reduce costs and doses, and improve the immunogenicity, of all major classes of SARS-CoV-2 vaccines.

## INTRODUCTION

Coronaviruses are enveloped single-stranded, positive-strand RNA viruses of the family *Corornaviridae* (*1*). At least seven coronaviruses infect humans: the α-coronaviruses HCoV-229E and HCoV-OC43, and the β-coronaviruses SARS-CoV (SARS-CoV-1), HCoV-NL63, CoV-HKU1, MERS-CoV, and the recently described SARS-CoV-2, a β-coronavirus closely related to human SARS-CoV-1 (79.0% nucleotide identity) and to SARS-CoV-like variants isolated from bats (*2–4*). SARS-CoV-2 infection causes flu-like symptoms in many patients, but in other cases develops into an acute pulmonary syndrome (*3, 5*). SARS-CoV-1 causes severe acute respiratory syndrome (SARS), whereas disease associated with SARS-CoV-2 has been named COVID-19. SARS-CoV-2, like SARS-CoV-1, requires expression of the cellular receptor ACE2 to infect cells (*6–8*).

Entry of SARS-CoV-2 into ACE2-expressing cells is mediated by its spike (S) protein (*7, 8*). The coronavirus S protein is a type I viral entry protein similar to influenza hemagglutinin and the HIV-1 envelope glycoprotein (*9*). Like these latter entry proteins, the S protein is processed into two domains, S1 and S2 (*7*). S1 binds ACE2, whereas S2 anchors the S protein to the viral membrane. The SARS-CoV-2 S protein has an efficient furin cleavage site at its S1/S2 boundary, and this site is processed in virus-producing cells (*10*). In contrast, the SARS-CoV-1 S1/S2 junction is cleaved by extracellular or target-cell proteases including TMPRSS2 and cathepsin L (*11–13*). Both S proteins require processing at a second site, S2’, within the S2 domain to mediate fusion of the viral and target cell membranes (*14*).

The receptor-binding domains (RBDs, also described as S^B^) of SARS-CoV-1 and SARS-CoV-2 directly bind ACE2 (*7, 15–17*). These RBDs are structurally and functionally distinct from the remainder of the S1 domain, and express and fold as independent domains (*15*). Both RBDs are highly stable and held together by four disulfide bonds. Structural studies of the SARS-CoV-2 RBD bound to ACE2 have identified a variable region, termed the receptor-binding motif (RBM), which directly engages ACE2 (*16*). This region is divergent between SARS-CoV-1 and SARS-CoV-2, although both RBD bind ACE2 in the same orientation and rely on conserved, mostly aromatic, residues to engage this receptor. The divergence between the SARS-CoV-1 and SARS-CoV-2 RBM domains suggest that this region is subject to ongoing positive selection from the humoral response in various hosts. However, some ten months into the pandemic, changes in the SARS-CoV-2 RBD remain exceedingly rare, consistent with a relatively slow overall rate of viral mutation throughout the genome.

Because the S protein is the major protein exposed on the virion, and because its activity can be impeded with antibodies, it is likely the major target of any SARS-CoV-2 vaccine. Soluble trimeric S proteins, including those stabilized through various mechanisms, have been tested as SARS-CoV-1 vaccines, and similar approaches are now being taken against SARS-CoV-2 (*7, 17, 18*). In fact, all of the vaccines likely to be available in the first half of 2021 express or deliver a full-length or ectodomain S protein, typically engineered with a pair of prolines designed to enhance the stability of these constructs (*19*). Nonetheless, the neutralizing activity of these vaccines correlates with RBD recognition, and the vast majority of potent neutralizing antibodies described to date, including those in late-stage clinical trials, target the RBD (*20–24*).

A different approach, immunizing with the RBD alone, has been shown to raise potent neutralizing antibodies against SARS-CoV-1 in rodents (*25, 26*). Although the RBD presents fewer epitopes than the S-protein trimer, this approach may have key advantages. First, a much larger fraction of RBD epitopes, essentially all RBD epitopes exposed on the native trimer, are neutralizing. Thus the RBD has fewer decoy epitopes and a greater fraction of the antibodies elicited will be neutralizing. Second, the 197-amino-acid RBD (S-protein residues 331-527) is much easier to produce than the full S-protein trimer. Thus the costs of production of a subunit vaccine will be lower, and expression from an mRNA or adenoviral vaccine will be greater, allowing dose sparing and limiting side effects. Third, an RBD-based vaccine is less likely to include linear or conformational epitopes that, in rare cases, might promote autoimmune disorders through molecular mimicry. Similarly, fewer epitopes reduce residual concerns about antibody-dependent enhancement, observed with other coronaviruses and primarily mediated through non-neutralizing epitopes. Finally, multivalent antigens are typically more immunogenic than dimeric or trimeric vaccines, and multivalency is much more easily obtained with RBD-based vaccines compared with those based on S-protein trimers.

Here we show that when equal amounts of the SARS-CoV-2 S-protein ectodomain or the RBD alone were conjugated to each of two carrier proteins, the RBD generated neutralizing responses equal to or greater than those of the S-protein. We nonetheless noted that the RBD expressed inefficiently, especially as a fusion protein with a range of multivalent carrier proteins. We therefore engineered the RBD to correct this deficiency and showed that this modified RBD, fused to five different multivalent carrier proteins and expressed as DNA vaccines, elicited a more potent neutralizing antibody responses than the wild-type RBD or the full-length proline-stabilized S-protein antigen used in several prominent COVID-19 vaccines. Finally we show that our modified RBD is more inherently immunogenic than the wild-type RBD when administered at equal dosage as a subunit protein vaccine. These data suggest that future vaccines against COVID-19 should include multivalent forms of engineered RBD antigens.

## RESULTS

### The SARS-CoV-2 RBD can elicit potent neutralizing antisera

The SARS-CoV-2 RBD, like that of SARS-CoV-1, is exposed in both known states of the S-protein trimer, namely a closed state where each RBD contacts symmetrically its analogues on the other protomer, and an open state in which at least one RBD domain is extended to contact ACE2. We have previously shown that the SARS-CoV-1 RBD folds independently and expresses efficiently, and that an immunoadhesin form of this RBD bound ACE2 more efficiently than constructs based on the S1 domain (*15*). This construct, RBD-Fc, also efficiently raised antibodies in mice capable of neutralizing SARS-CoV-1 variants, including those with distinct RBD sequences (*25, 26*). Moreover, the vast majority of well characterized neutralizing antibodies against SARS-CoV-2, including those in late-stage clinical trials, target the RBD. These data suggest that an RBD-based SARS-CoV-2 vaccine could be effective against virus throughout the current COVID-19 pandemic.

To initially evaluate this possibility under optimal conditions, we evaluated the immunogenicity of the SARS-CoV-2 RBD fused to the Fc domain as an expedient for rapid purification. RBD-Fc was chemically conjugated to a Keyhole limpet hemocyanin (KLH) carrier protein and mixed with the AS01 adjuvant formulation now used in at least two human vaccines (**Fig. 1**). This antigen/adjuvant combination was inoculated intramuscularly into four female Sprague-Dawley rats with a schedule of seven increasing (2.5-fold) doses, one each day, ultimately administering a total of 500 μg of the SARS-CoV-2 RBD-Fc. Thirty days after the first administration, RBD fused to a 4-amino acid C-tag was purified with a C-tag affinity column and administered as before. Blood was harvested from each of the four rats (R15, R16, R17, R18) immediately before inoculation (day 0) and 40 days after the first inoculation. Serial dilutions of day-0 and day-40 sera were measured for their ability to neutralize retroviruses pseudotyped with the SARS-CoV-2 S protein (SARS2-PV). To estimate neutralization potency, these sera were also compared with a mixture of all four day-0 preimmune sera, further combined with an immunoadhesin form of ACE2 (ACE2-Fc) at concentrations of 10, 100, and 1000 μg/ml before dilution. As anticipated, some baseline inhibition could be observed in heat-inactivated rat sera (grey lines in **Fig. 1A**). However day-40 serum from each rat, obtained after two sets of immunizations, potently neutralized SARS2-PV entry with an efficiency comparable to or greater than day-0 preimmune sera mixed with ACE2-Fc at a 100 μg/ml concentration (**Fig. 1, A and B**). We conclude that the SARS-CoV-2 RBD can elicit a neutralizing response in vaccinated rats comparable to a 100 μg/ml (1 μM) concentration of an inhibitor with a 1 nM IC_50_. To confirm that sera from vaccinated rats neutralized SARS2-PV by recognizing the SARS-CoV-2 S protein, we used each pooled serum to prevent binding of an ACE2-Fc variant bearing a rabbit Fc domain (ACE2-rIg) from cells expressing the SARS-CoV-2 S protein (**Fig. 1C**). The ability of pooled serum to compete with ACE2-rIg indicates that these antisera neutralized SARS2-PV entry by blockading ACE2 association with the S protein. Thus, under ideal conditions, immunization with SARS-CoV-2 RBD elicits antibodies that very potently neutralize SARS2-PV, and these antibodies do so by preventing S-protein association with ACE2.

**Fig. 1.**
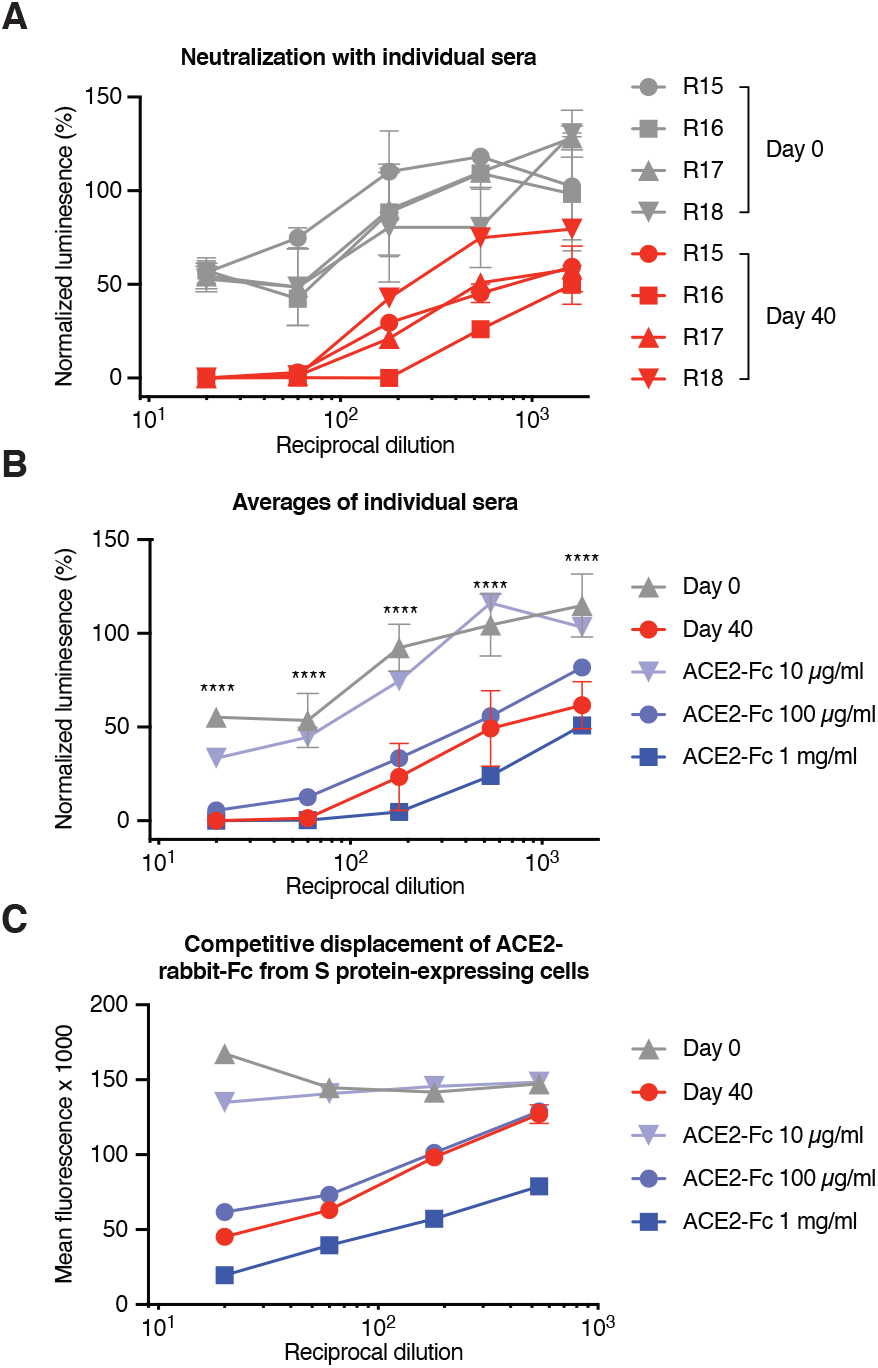
Immunization with the SARS-CoV-2 RBD elicits potently neutralizing antibodies. Four female Sprague Dawley rats (R15, R16, R17, R18) were immunized with two sets of escalating doses of RBD conjugated to keyhole limpet hemocyanin. (**A**) The indicated dilutions of preimmune sera (day 0, gray) were compared to dilutions of sera harvested from immunized rats at day 40, and to the same dilutions of preimmune sera mixed to achieve the indicated ACE2-Fc concentrations before dilution. Each serum and serum-ACE2-Fc mixture was compared for its ability to neutralize S-protein-pseudotyped retroviruses (SARS2-PV), by measuring the activity of a firefly-luciferease reporter expressed by these pseudoviruses. Figure shows entry of SARS2-PV as a percentage of that observed without added rat serum. Error bars indicate the range of two neutralization studies. (**B**) The data from each rat of panel A are averaged for clarity. Error bars indicate s.d., with each rat considered a different experiment. Differences between day-0 and day-40 serum are significant at all dilutions (*P* < 0.001; two-way ANOVA). (**C**) Pooled sera and pooled preimmune sera mixed with the indicated concentrations of ACE2-Fc were further combined with an ACE2-Fc variant bearing a rabbit-derived Fc domain. Binding of the ACE2-Fc was monitored with an anti-rabbit Fc secondary antibody as determined by flow cytometry. Error bars indicate the range of two such measurements. Differences between day-0 and day-40 serum are significant (P < 0.001; two-way ANOVA) at all dilutions.

One concern associated with coronavirus vaccines is the possibility that anti-S-protein antibodies could promote infection of cells, such as alveolar macrophages, expressing Fc receptors, for example FcγRI (CD64) or FcγRII (CD32). This undesirable antibody-dependent enhancement (ADE) has been well characterized in tissue-culture studies of several flaviviruses including Zika virus (ZIKV) and dengue virus (*27*). To evaluate this possibility for SARS-CoV-2, SARS2-PV were mixed with pooled day-0 or day-40 serum at the indicated serial dilutions, and the resulting virus/sera mixtures were incubated with HEK293T cells transfected to express rat FcγRI. These cells did not express ACE2 and no infection was observed with day-0 preimmune sera nor day-40 immune sera (**Fig. S1A**). In contrast, rat anti-ZIKV antisera, or day-0 preimmune sera incubated at the same dilutions with ZIKV virus-like particles (VLP) promoted robust ADE (**Fig. S1B**). ADE activity peaked at approximately a 3000-fold dilution, consistent with competition between ADE and neutralizing activities of these antisera. Moreover, no SARS2-PV ADE was observed in the presence of ACE2 (**Fig. S2, C and D**) or with K562 cells that endogenously express FcγRII (**Fig. S1, E and F**) (*28*). Thus anti-RBD antisera do not mediate SARS2-PV ADE in the presence or absence of ACE2 under the described conditions.

### The RBD is more immunogenic than the S-protein ectodomain

To directly compare the immunogenicity of the RBD and a proline-stabilized S-protein ectodomain, both proteins were fused at their C-termini to a SpyTag and conjugated to one of two carrier systems (**Fig. 2**). First, each was conjugated by isopeptide bond formation to SpyCatcher-mi3 60-mer particles (RBD-mi3 and S-mi3) (*29, 30*). Second, the same proteins were conjugated chemically to KLH as in Figure 1 (RBD-KLH and S-KLH). Four rats were vaccinated with each antigen/carrier combination as in Fig. 1 except that one-fifth the total antigen (100 μg) was administered for each vaccination round. Sera collected at day 0 and day 45 from each rat was characterized for neutralization with SARS2-PV (**Fig. S2**) and the results were averaged (**Fig. 2, A and B**). We observed that, with both carrier proteins, RBD conjugates elicited more potent neutralizing responses at day 45 than did S-protein conjugates. We further observed that conjugates to the mi3 60-mer elicited more potent responses than conjugates to the KLH carrier protein. Thus RBD-mi3 was significantly more immunogenic than S-mi3, RBD-KLH and S-KLH (**Fig. 2C**). We conclude that, when equal amounts of the RBD and the S-protein ectodomain are conjugated to a carrier and administered with a potent adjuvant, the RBD elicits a more potent neutralizing response. We also conclude that the mi3 carrier protein elicited more potent responses to both antigens than did KLH.

**Fig. 2.**
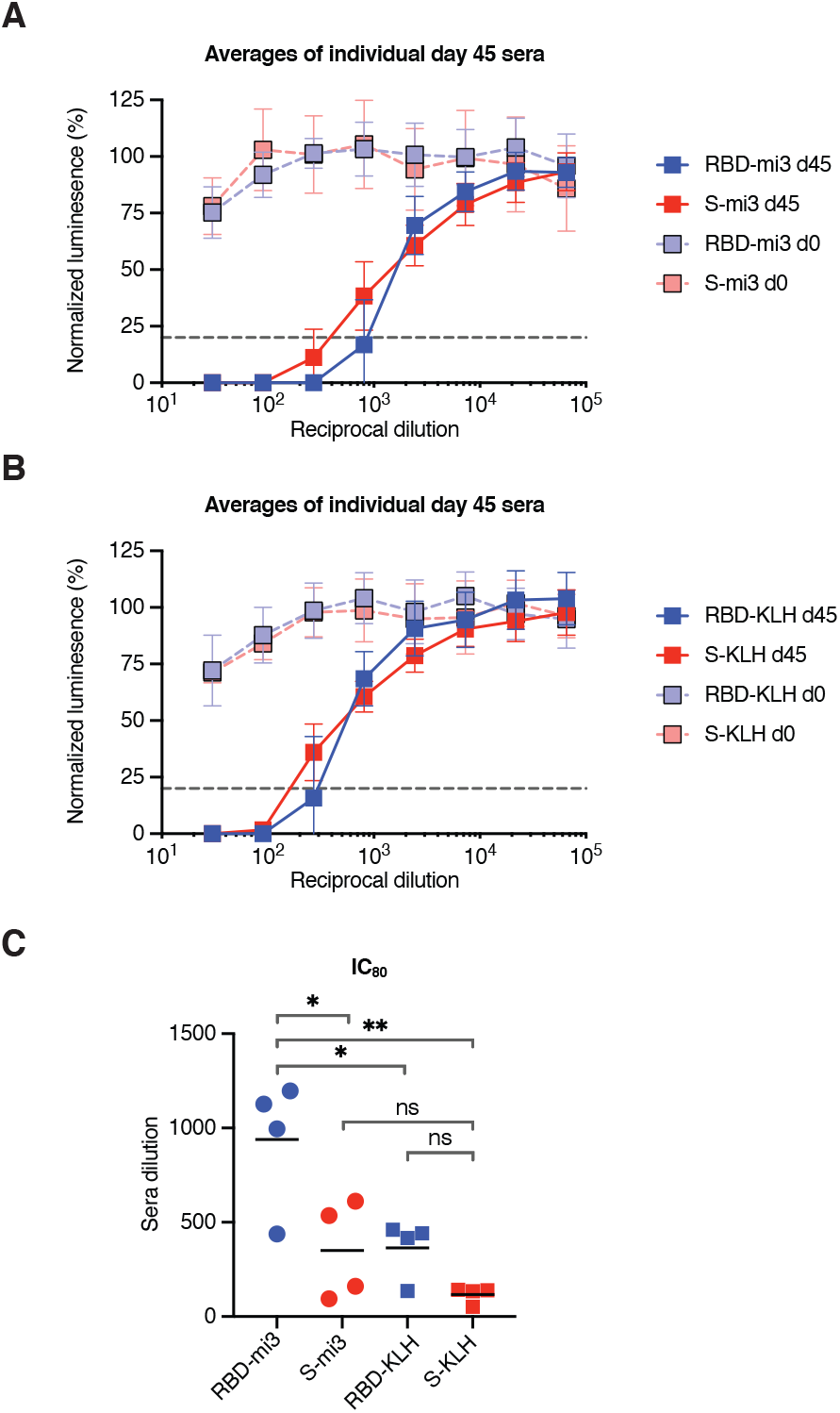
SARS-CoV-2 RBD nanoparticles are more immunogenic than S-protein nanoparticles. Four groups of four female Sprague Dawley rats were inoculated with either RBD-SpyTag or S-protein-SpyTag conguated to either SpyCatcher-mi3 particles (**A**) by isopeptide bond formation, or KLH (**B**) by EDC. The indicated dilutions of preimmune sera (day 0) were compared to dilutions of sera harvested from immunized rats at day 40. Each serum was compared for its ability to neutralize S-protein-pseudotyped retroviruses (SARS2-PV), by measuring the activity of a firefly-luciferease reporter expressed by these pseudoviruses. Figure shows entry of SARS2-PV as a percentage of that observed without added rat serum. Dotted lines indicate 80% neutralization. Error bars indicate s.d. for biological replicates. (**C**) IC_80_ values for each rat at day 40 were calculated in Prism 8 and significance between groups is indicated (* indicates *P* < 0.05; ** indicates *P* < 0.01; ns indicates *P* > 0.05; one-way ANOVA with Tukey’s multiple comparison test).

### A glycan-modified RBD, gRBD, administered as a protein, elicits a more potent neutralizing response than does the wild-type RBD

Conjugates of the sort produced for Fig. 2 cannot readily be used with genetic vaccines such as those delivered as DNA, or as mRNA, or through viral vectors. However fusion proteins that express both the antigen and the carrier as a single polypeptide chain can be used in these formats. Such fusion proteins also simplify the production of subunit protein vaccines. We therefore undertook to produce fusion proteins with the RBD in various formats but observed that these constructs were not efficiently produced in cells and were not efficiently secreted (**Fig. S4**). To solve this problem, we developed a screening procedure whereby the expression levels of engineered RBD variants, expressed as dimers, were monitored by flow cytometry. We observed that one such variant, modified with four additional glycosylation sites in an RBD region occluded in S-protein trimer (**Fig. 3, A to C**), expressed and secreted markedly more efficiently as a fusion with the mi3 60-mer than did the unmodified RBD (**Fig. 4A**). Each of three of these newly engineered glycans – those engineered at residues 370, 428, and 517 – markedly increased RBD expression when fused to a multivalent carrier (**Fig. S3**), with the engineered glycosylation motif at residue 517 contributing most to gRBD expression. A fourth glycan, at residue 394, did not contribute to higher RBD expression, but it was included to further limit RBD aggregation.

**Fig. 3.**
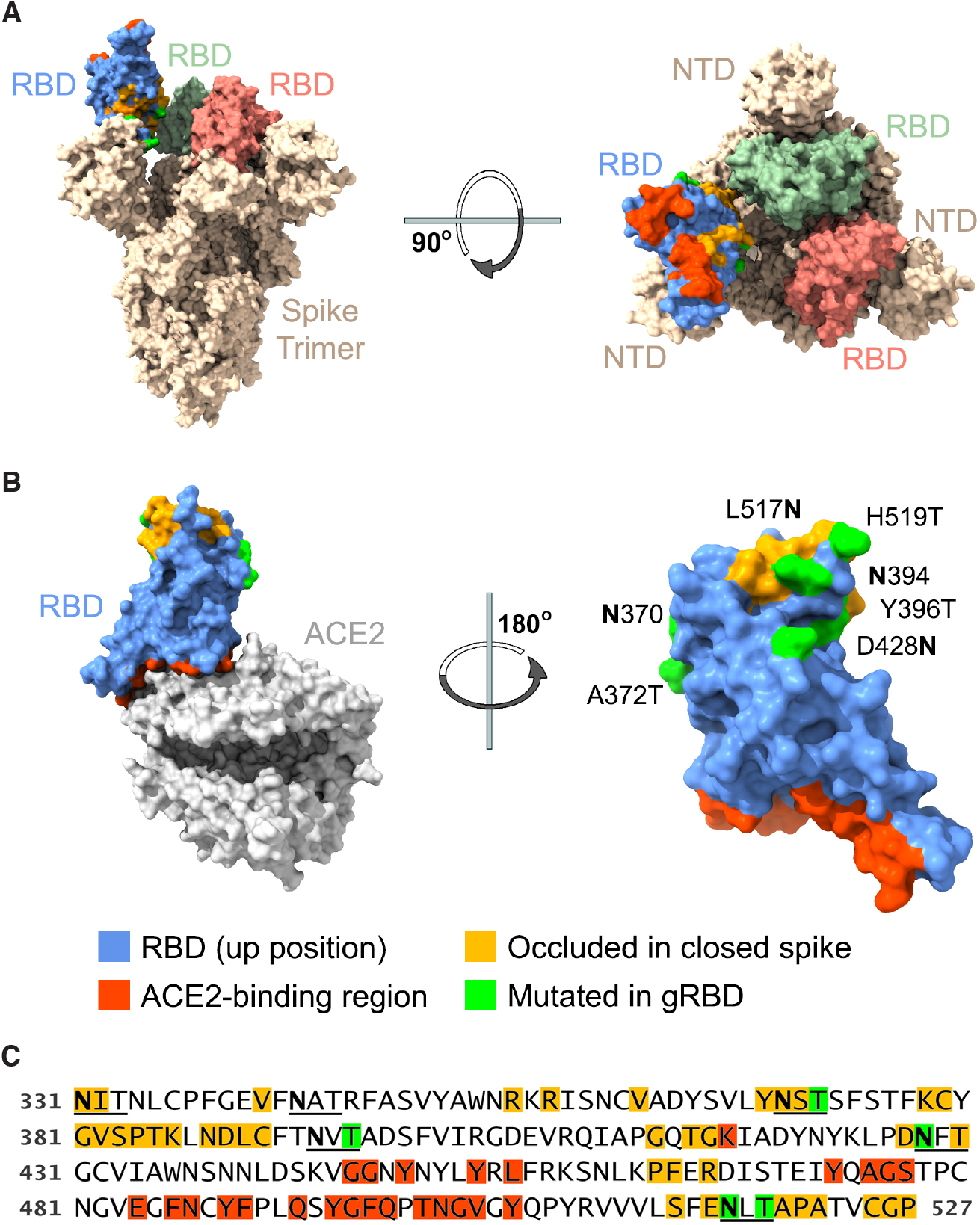
Engineered SARS-CoV-2 RBD glycans enhance expression of multivalent RBD fusion proteins. Views of the RBD (**A**) in the context of the SARS-CoV-2 S protein in the open one-up conformation, with the ACE2-binding region (red) facing upward and (**B**) bound to the ACE2 receptor, with the RBD ACE2-binding region facing downwards. Blue indicates surface residues that are neither occluded in the closed conformation (indicated by yellow) nor part of the ACE2 interface (red). Green indicates residues whose mutation creates a novel N-glycoslylation motif. (**C**) The sequence of the engineered RBD bearing four novel glycosylation motifs (gRBD) is shown. Numbering indicates S-protein residue. Glycosylation motifs (2 native and 4 engineered) are underlined. Coloring is as described in (B).

**Fig. 4.**
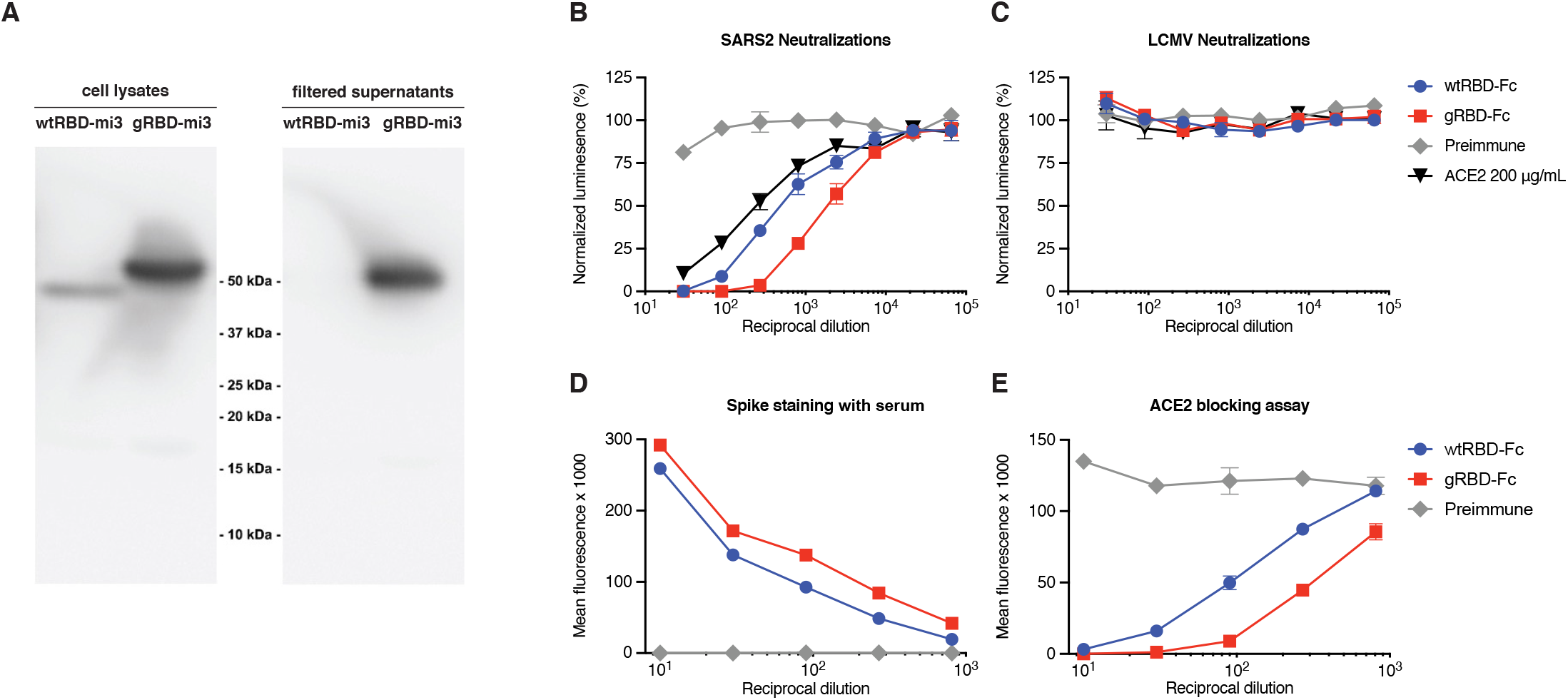
gRBD expresses efficiently as an mi3 fusion protein and is more immunogenic than wild-type RBD as adjuvanted protein. Expression: (**A**) RBD-mi3 60-mer fusion proteins were expressed in Expi293 cells, after 5 days, supernatants and cell lysates were analyzed by SDS-PAGE anti-CTag western blot. Note that no wtRBD-mi3 could be detected in the supernatant. Immunogenicity: Five mice per group were inoculated with 25 μg of protein A/SEC purified wtRBD-Fc or gRBD-Fc adjuvanted with 25 μg of MPLA and 10 μg Quil-A. Immunizations were conducted day 0 and day 14, and serum was collected and pooled on day 21. Pooled preimmune sera, and pooled preimmune sera doped with 200 μg/mL of ACE2-Fc were used as negative and positive controls. Pooled sera were used to neutralize (**B**) SARS-CoV2 pseudovirus or (**C**) LCMV pseudovirus control. In parallel, HEK293T cells were transfected with 1 μg / well in a six well plate and stained the next day with pooled preimmune and day-21 sera and then stained with either (**D**) anti-mouse-FITC or (**E**) ACE2-Fc-DyLight650. Error bars indicate SEM.

We then investigated whether these additional glycans would alter immune responses to the RBD (**Fig. 4, B to E**). Because fusions of the wild-type RBD with higher-order multivalent carrier proteins proved difficult to express and purify, we produced RBD and gRBD as fusion proteins with Fc domains of human IgG1 (wtRBD-Fc and gRBD-Fc). Two doses of 25 μg of RBD-Fc antigen with 25 μg MPLA and 10 μg QuilA adjuvants, separated by 14 days, were administered intramuscularly to five mice per group. Anti-sera were harvested 21 days after the first vaccination, and analyzed for its ability to neutralize SARS2-PV or control pseudoviruses expressing the entry (GP) protein of the lymphocytic choriomengingitis virus (LCMV-PV). Sera from inoculated mice was mixed and compared with pre-immune sera and pre-immune sera mixed at an initial concentration of 200 μg/ml ACE2-Fc. We observed, somewhat unexpectedly, that gRBD-Fc elicited a more potent neutralizing response than did wtRBD-Fc (**Fig. 4, B and C**). Consistent with this observation, sera from gRBD-Fc inoculated mice more efficiently bound cell-expressed S protein (**Fig. 4D**) and more efficiently blocked binding of fluorescently labeled ACE2-Fc (**Fig. 4E**) than did sera from wtRBD-Fc-inoculated mice. Thus the engineered glycans of gRBD do not interfere with, and may enhance its ability to raise anti-RBD antibodies in mice. We speculate that gRBD glycans better focus the B-cell response to neutralizing RBD epitopes (*31, 32*). Alternatively, aggregation of the wild-type RBD may impede access to these epitopes. Collectively these data suggest that multivalent antigens based on gRBD will be easier to produce and at least as immunogenic as their wild-type RBD analogues.

### DNA vaccines expressing multivalent gRBD fusion proteins elicit more potent neutralizing antisera than do the corresponding wtRBD fusion proteins or the full-length S protein

To evaluate the utility of gRBD in the context of DNA-, mRNA-, or viral vector-based vaccines we developed plasmids expressing wtRBD and gRBD-fusion proteins with five multivalent carriers (**Fig. 5**). Specifically, fusions with the IgG1 Fc-domain dimer (Fc), the T4 foldon trimerizing domain (*33*), a dodecameric scaffold based on the *H. pylori* neutrophil-activating protein (NAP) (*34*), the *H. pylori* ferritin 24-mer (*35*), and the engineered mi3 60-mer (*29, 30*). Complete amino-acid sequences of these fusion proteins are provided in **Fig. S4A**. In each case, the gRBD fusion protein expressed more efficiently than its wild-type RBD analogue (**Fig. S4B**). We also evaluated a plasmid expressing the full-length proline-stabilized 1273 amino-acid S protein (S1273-PP, the full-length S protein with prolines introduced at residues 986 and 987) which expressed efficiently on the surface of HEK293T cells, more so than the otherwise identical construct lacking these stabilizing prolines. (**Fig. S4C**). Five mice per group were electroporated with 120 μg plasmid (60 μg per hind leg) encoding either wtRBD or gRBD antigen fused to each of the aforementioned multivalent scaffolds, or with plasmid expressing the S1273-PP full-length S protein. Mice were again electroporated 14 days later with the same plasmds. Sera were harvested 21 days after the first inoculation. Combined sera for each group were evaluated for its ability to neutralize SARS2-PV or LCMV-PV and compared with preimmune sera or preimmune sera mixed with ACE2-Fc at an initial concentration of 200 μg/ml. In each case, sera from gRBD-fusion constructs neutralized SARS2-PV more efficiently than their wtRBD analogues, and more efficiently than sera from mice electroporated with S1273-PP (**Fig. 5A**). Among the various scaffolds, the *H. pylori* ferritin 24-mer elicited the most potent neutralizing antisera (**Fig. 5, B and C**). We conclude that gRBD consistently and significantly (*P*=0.0089, **Fig. 5C**) improves the immunogenicity of multivalent fusion proteins relative to the same construct fused to the wild-type RBD.

**Fig. 5.**
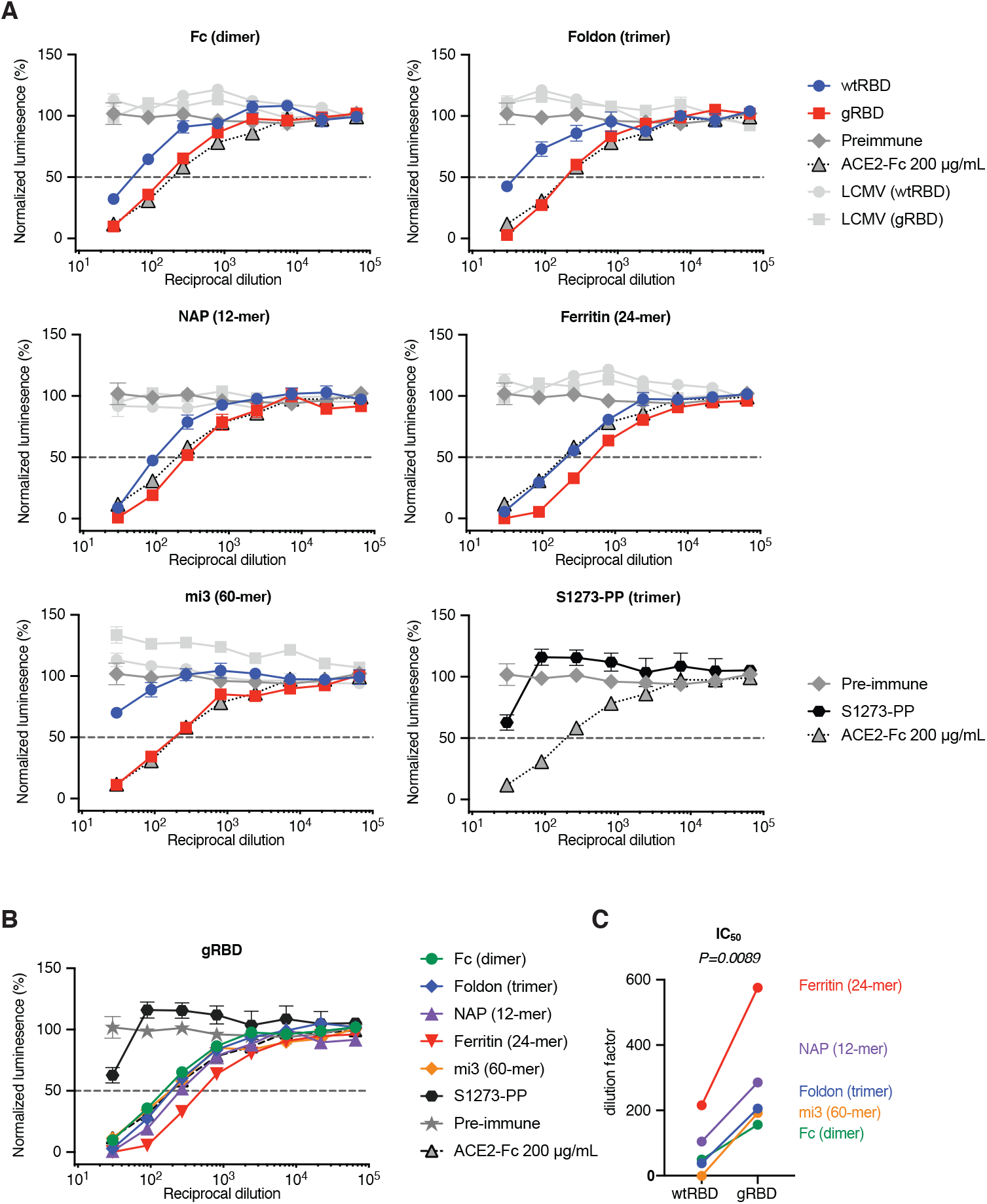
gRBD based DNA vaccines more efficiently raise neutralizing antibodies than those based on wild-type RBD. **(A)** Five mice per group were electroporated in each hind leg with 60 μg plasmid DNA expressing wtRBD or gRBD fused to human Fc dimer, foldon trimer, *H. pylori* NAP 12-mer, *H. pylori* ferritin 24-mer, and mi3 60-mer. An additional control group was electroporated with plasmid expressing the full-length SARS-CoV-2 spike protein with two stabilizing prolines (S1273-PP). Electroporations were conducted day 0 and day 14, and serum was collected and pooled for neutralization assays on day 21. Pooled preimmune sera, and pooled preimmune sera doped with 200 μg/mL of ACE2-Fc were used as negative and positive controls. (**B**) Neutralizing potency of gRBD varied by platform. (**C**) IC_50_ calculations for wtRBD and gRBD were calculated (Prism 8) against normalized values by least squares fit. P-value was calculated by 2-tailed paired t test between wtRBD and gRBD pairs.

## DISCUSSION

For several reasons, a vaccine against SARS-CoV-2 should be easier to develop than those against many other viruses (*9, 36*). First, coronaviruses have exceptionally large genomes compared to other RNA viruses, and, to avoid error catastrophe, their viral polymerase has acquired a proof-reading function. Thus any individual gene, for example that of the S protein, is likely to retain its original sequence over multiple replication cycles. Indeed now 9 months into the SARS-CoV-2 pandemic, few mutations in the S protein, and no dominant mutations in its RBD, have yet been described. Second, coronaviruses in general, and clearly SARS-CoV-2 in particular, transmits to a new host more rapidly than an adaptive immune response can emerge. A likely consequence of this strategy is that one of its most critical epitopes, namely its RBD, is exposed on the virion (*7, 9, 15, 17*), favoring transmission efficiency over antibody resistance. Finally, the stability and compactness of the SARS-CoV-1 and SARS-CoV-2 RBD suggests that it can be easily manufactured and presented to the immune system using many production technologies, presentation scaffolds, and delivery systems (*15*). Here we repeat studies of the SARS-CoV-1 RBD showing that the SARS-CoV-2 RBD is sufficient to raise potent neutralizing antibodies (**Fig. 1**), and that it does so more efficiently than the SARS-CoV-2 S protein ectodomain when conjugated to a KLH or mi3 carrier and administered as a protein (**Fig. 2**).

However, the wild-type SARS-CoV-2 RBD suffers from one major deficiency. When expressed as a fusion protein in multivalent scaffold or carrier, it is expressed inefficiently in cells, it secretes poorly, and it tends to aggregate (**Fig. 4A**, **Fig. S4B**). This tendency to aggregate appears to impair its immunogenicity, even when administered as a reasonably well-behaved Fc dimer (**Fig. 4B**). Through trial-and-error screening of RBD variants modified in a region normally occluded on the S-protein trimer, we identified a construct with four novel glycosylation motifs (**Fig 3**), that substantially improved or rescued expression of all five multivalent protein carriers assayed (**Fig. S4B**). Each of these glycosylation motifs improved expression or RBD solubility (**Fig. S3**). The resulting engineered RBD, which we call gRBD to reflect its additional glycans, elicited more potent neutralizing sera when administered as an adjuvanted protein (**Fig. 4, B to E**) or when electroporated as a DNA vaccine expressing each of five carrier proteins (**Fig. 5, A** to **C**). Importantly, these gRBD fusion proteins were more immunogenic than proline-stabilized S proteins used as antigens in most prominent SARS-CoV-2 vaccines (**Fig. 5B**).

Our data therefore show that (1) the SARS-CoV-2 RBD can be more immunogenic than the S protein, (2) an RBD engineered with four glycans can be more immunogenic than the wild-type RBD, and (3) multivalent forms of the RBD and gRBD, and in particular those fused with the *H.pylori* ferritin 24-mer, can be more immunogenic than dimeric or trimeric constructs. Why is the RBD more immunogenic than the S protein? The answer appears straightforward: the RBD is the dominant neutralizing epitope. Expression of the remainder of the protein taxes cellular resources and exposes potential decoy epitopes. Why is gRBD more immunogenic than the wild-type RBD? We speculate that poor folding or solubility of the wild-type RBD help occlude its major neutralizing epitopes or limit its access to the lymph nodes (*37*). It is also possible that the glycans of gRBD mask dominant but non-neutralizing RBD epitopes. Finally, why is the ferritin 24-mer more immunogenic than the other scaffolds? Immunogenicity is likely improved by multivalency, by higher expression, and by pre-existing T cell help (*37, 38*). The mi3 60-mer is maximally multivalent, but it expresses relatively poorly and may include fewer epitopes recognized by established memory T cells. The NAP 12-mer expresses much more efficiently, but its size or the arrangement of gRBD domains may be sub-optimal. The Fc dimer and foldon trimer may also be insufficiently multivalent, and again include fewer T-cell epitopes. The *H. pylori* ferritin 24-mer combines high expression, a larger particle and high valency, and it provides T-cell epitopes similar to those in many bacterial antigens. Further work identifying and engineering high-expressing multivalent scaffolds capable of presenting gRBD may yield even more potent antigens.

In short, we have engineered a SARS-CoV-2 RBD antigen that expresses more efficiently than the wild-type RBD as a fusion with multivalent carrier proteins, and these fusion proteins are more immunogenic as protein or DNA vaccines than commonly used S protein antigens. We propose therefore that multivalent gRBD fusion proteins could improve production efficiencies of protein-based SARS-CoV-2 vaccines, and limit the doses necessary for all vaccine classes.

## MATERIALS AND METHODS

### Production of SARS-CoV-2 and LCMV pseudoviruses and Zika-virus virus-like particles

Retroviruses pseudotyped with the SARS-CoV-2 S protein or lymphocytic choriomenginitis virus GP protein (SARS2-PV, LCMV-PV) were produced as previously described (Moore et al., 2004) with modest modifications as described. HEK293T cells were transfected by polyethylenimine (PEI) transfection at a ratio of 5:5:1 with a plasmid encoding murine leukemia virus (MLV) gag/pol proteins, a retroviral vector pQCXIX expressing firefly luciferase, and a plasmid expressing the spike protein of SARS-CoV-2 (GenBank YP_009724390) or LCMV GP (GenBank AHZ55917.1). Cells were washed 6 hours later, and the culture supernatant containing pseudoviruses was harvested at 48-72 hours post transfection. Zika virus virus-like particles (ZIKV-VLP) were produced by transfecting HEK293T cells by the calcium phosphate transfection method with a ZIKV replicon (strain FS13025, GenBank KU955593.1), whose expression is controlled by tetracycline, a plasmid encoding ZIKV capsid, prM, and E proteins (strain FSS13025, GenBank KU955593.1), and the pTet-On plasmid expressing a reverse Tet-responsive transcriptional activator (rtTA) at a ratio of 2:1:1. Cells were washed 6 hours later and replenished with fresh media containing 1 μg/ml doxycycline. The VLP-containing culture supernatant was harvested 48 h post transfection. ZIKV replicon was generated by replacing the region spanning 39^th^ through 763^rd^ amino acids of the polyprotein of a ZIKV molecular clone we previously generated (*39*) with Renilla luciferase with the 2A self-cleaving peptide fused atits C-terminus. This construct contains the tetracycline-responsive Ptight promoter that drives ZIKV RNA transcription. The pseudovirus- or VLP-containing culture supernatants were cleared by 0.45-μm filtration. SARS2-PV and ZIKV-VLP titers were assessed by RT-qPCR targeting the CMV promoter in the retroviral vector pQCXIX and ZIKV NS3 gene, respectively. In some cases, clarified pseudovirus and VLP stocks were stored at −80°C for long-term storage and reuse.

### Generation of human ACE2 expressing cell lines

HEK293T cells expressing human ACE2 (hACE2) were generated by transduction with murine leukemia virus (MLV) pseudotyped with the vesicular stomatitis virus G protein and expressing myc-hACE2-c9, as previously described (Wicht et al., 2014). Briefly, HEK293T cells were co-transfected by PEI with three plasmids, pMLV-gag-pol, pCAGGS-VSV-G and pQCXIP-myc-hACE2-c9 at a ratio of 3:2:1, and medium was refreshed after overnight incubation of transfection mix. The supernatant with produced virus was harvested 72-hours post transfection and clarified by passing through 0.45μm filter. 293T-hACE2 cells transduced with MLV vectors were selected and maintained with medium containing puromycin (Sigma). hACE2 expression was confirmed by SARS1-PV and SARS2-PV entry assays and by immunofluorescence staining using mouse monoclonal antibody recognizing c-Myc.

### Protein Production

Expi293 cells (Thermo-Fisher) were transiently transfected using FectoPRO (Polyplus) with plasmids encoding SARS-CoV2 RBD with a human or rabbit-Fc fusion or a C-terminal C-tag (-EPEA). After 5 days in shaker culture, media were collected and cleared of debris for 10 min at 3,000 g and filtered using 0.45-μm flasks (Nalgene). Proteins were isolated using MabSelect SuRe (GE Lifesciences) or CaptureSelect C-TagXL (Thermo-Fisher) columns according to the manufacturers’ instructions. Eluates were buffer exchanged four times with PBS and concentrated using Amicon ultra filtration devices (Millipore Sigma), except for S-protein-Spytag, which was buffer exchanged by dialysis (Pierce) 3 times, and concentrated. For wtRBD-Fc and gRBD-Fc used in mouse protein immunizations, further purification was performed by SEC on a HiPrep 16/60 Sephacryl S-400 HR column connected to an ÄKTA FPLC. Fractions were isolated with PBS buffer, verified by SDS-PAGE gels, pooled, and concentrated. All purified proteins were stored at 4°C prior to use.

For RBD-multimer fusion proteins, supernatant pH was adjusted to 8.5 by addition of 1/20 volume 1M Tris pH=9.0 (G-Biosciences, 786-476), and the supernatant was filtered by 0.45 μm filter prior to purification on CaptureSelect™ C-tagXL Columns. In the case of RBD-ferritin fusions, 0.5% Tween-20 was also added to supernatants prior to filtration, and supernatants were mixed with anti-flag M2 agarose affinity gel (1 mL slurry for 25mL culture, Sigma-Aldrich, A2220-10ML), and incubated on a rotary shaker overnight at 4°C. The mixture was packed into columns (Agela Technologies, AZ-IC-1T). Columns (C-tag or M2) were washed with 10 CV TBS (25 mM Tris 8.5, 150 mM NaCl), and eluted with 5 CV Gentle Ag/Ab Elution Buffer, pH 6.6 (Thermo Scientific Pierce, 21027). Buffer was exchanged 4 times with TBS 8.5 for yield studies, or 3 times with TBS 8.5, and one time with PBS for subsequent SEC polishing.

### Protein immunizations and sera collection in rats

All animals used in these studies were handled and maintained in accordance with NIH guidelines and approved by Institutional Animal Care and Use Committee (IACUC) of Scripps Research (Protocol 18-025). Female Sprague Dawley rats were immunized with incremental increasing doses of antigen over seven days starting at day 0, and boosted with a similar regimen at day 30.

Rats were inoculated in the first set with the SARS-CoV-2 RBD fused to the Fc domain of human IgG1, and in the second set with the RBD fused to a four amino-acid C-tag. In both cases RBD fusions were conjugated at a 1:1 ratio to Mariculture keyhole limpet hemocyanin (mcKLH, Thermo-Fischer Peirce) by 1-ethyl-3-(3-dimethylaminopropyl)carbodiimide hydrochloride (EDC, Thermo-Fischer Peirce) according to manufactures protocols. Each set of seven injections were performed in the following manner. RBD-KLH conjugates were administered intramuscularly into the rear quadriceps. Inoculations were initiated with 2.2 μg RBD-KLH (equivalent to 1.1 μg RBD antigen) adjuvanted with 0.1 μg MPLA and 0.1 μg QuilA, and this inoculum was increased by 2.55 fold for each of the next six days, administering a total of 500 μg RBD-Fc or RBD-C-tag fusion protein, and 40 μg of each adjuvant component. Sera were collected before inoculation (day 0 preimmune sera) and every five days starting on 10^th^ day after the first injection. All sera were heat-inactivated for 30 minutes at 56°C and stored at −80°C for reuse.

For the mi3 vs KLH round of inoculations, rats were inoculated with either RBD-Spytag or S-protein-Spytag, conjugated either to equal quantities of either Spycatcher-mi3 (mixed and incubated 4°C overnight) or mcKLH (EDC conjugation). Incremental increasing injections were conducted as above, but with an initial inoculation of 0.4 μg (0.2 μg each antigen and carrier) for a total of 200 μg protein over both rounds of vaccination and 40 μg of each adjuvant component.

### Protein immunizations and sera collection in mice

Female 8 to 9-week-old BALB/cJ mice were immunized with 25 μg protein antigen, 25 μg MPLA and 10 μg of QuilA on day 0 and day 14. Sera were collected before the initial inoculation (preimmune sera) and on day 21. All sera were heat-inactivated for 30 minutes at 56°C and stored at −80°C for reuse.

### DNA immunizations in mice

CMV/R expression plasmids encoding wtRBD or gRBD fused to multimerization platforms (Figure S3A) were prepared using NucleoBond® PC 2000 (Takara Bio USA Inc) and confirmed to be essentially endotoxin free using Pierce™ Chromogenic Endotoxin Quant (Thermoscientific) according to the manufacturers’ instructions. Female 8 to 9-week-old BALB/cJ mice were electroporated with 60 μg DNA in each hindquarter for a total dose of 120 μg on day 0 and day 14. Electroporations were conducted on a Harvard Apparatus ECM 839 BTX in LV mode at 40V using 8 pulses, pulse length of 100ms, at 100ms intervals with unipolar polarity. Sera were collected before the initial inoculation (preimmune sera) and on day 21. All sera were heat-inactivated for 30 minutes at 56°C and stored at −80°C for reuse.

### Neutralization studies of SARS-CoV-2 and LCMV pseudoviruses

Individual sera or pooled sera collected at day 0 (pre-immune sera), and at the indicated day after the first inoculation were serially diluted in DMEM. In some cases, day 0 sera were mixed with ACE2-Fc to a concentration of 10, 100, or 1000 μg/ml before dilution, and then diluted in the same manner. Individual, pooled, or pooled-ACE2-Fc sera was mixed with SARS2-PV and incubated at 37°C for one hour. For rat studies, one hour later, 104 ACE2-239T cells were added along with DEAE-Dextran (final concentration 5 μg/ml), and media was exchanged 6 hours later with fresh media without rat sera. For mouse studies, one hour later, 10^4^ ACE2-239T cells were added and spun at 3000×g for 30 minutes at 4°C, was then returned to 37°C, and media was exchanged 2 hours later with fresh media without mouse sera. At least two independently mixed replicates were measured for each experiment. Firefly luciferase activity was measured (Britelight) 48 hours post-infection. All neutralization studies were performed at least twice with similar results.

### Competitive displacement of ACE2-rabbit-Fc from S-protein expressing cells

Serial dilutions of pooled sera or pooled pre-immune sera mixed with ACE2-Fc (with the human Fc domain) at initial concentrations of 10, 100 and 1000 μg/mL were then mixed with 1 μg/ml ACE2-rabbit-Fc. Independent mixtures were made for each replicate. Pooled sera and sera mixtures were used to stain HEK293T cells transfected by the jetPRIME Transfection Reagent (Polyplus) to express the full-length SARS-CoV2 S protein. Specifically, 10^5^ per cells per well were placed in a 96-well V-bottom plate and incubated for 45 minutes at 4°C with 100 μl of the serially diluted sera doped with ACE2-Fc. After washing, cells were stained with anti-rabbit IgG-Alexa647 antibody for 45 minutes at 4°C, and mean fluorescence intensities were measured for each well by flow cytometry.

### Measurement of antibody-dependent enhancement

The ability of anti-SARS-CoV-2 RBD immune sera to mediate antibody-dependent enhancement (ADE) was measured using HEK293T cells or HEK293T cells stably expressing human ACE2 (293T-hACE2 cells), transfected using the calcium phosphate transfection method to express the rat ortholog of FcγRI (CD64). The human monocytic cell line K562 (ATCC CCL-243), which endogenously expresses FcγRII, was also used for ADE assays. The RBD immune sera, collected from four different rats at day 40 after the first immunization, were mixed at an equal ratio, as was preimmune sera obtained at day 0 from the same rats. As a positive control, sera from ZIKV-infected rats (rat #13 and #15, distinct from the similarly numbered RBD- inoculated rats), was also mixed at an equal ratio. Immune and pre-immune serum samples were heat inactivated for 30 min at 56°C and serially diluted in DMEM containing 10% heat-inactivated FBS. SARS2-PV or ZIKV-VLP in 50 μl was preincubated for 1 h at 37°C with 50 μl of diluted sera and added to the indicated cells plated on the 96 well plates. Two days later, infection levels were assessed using Luc-Pair Firefly Luciferase HS Assay Kit (Genocopia) for SARS-PV and Luc-Pair Renilla Luciferase HS Assay Kit (Genocopia) for ZIKV-VLP.

### SDS-PAGE Western Blotting

Expi293 supernatant samples were centrifuged and separated and filtered (0.45-μm). Equivalent fractions of cell lysate and clarified supernatants were loaded and run by SDS-PAGE on NuPAGE™ bis-tris gels. Gels were then transferred to 0.4-μm PVDF membrane using Xcell Blot Module under constant voltage of 25V for 1 hour. The membrane was wash and blocked (5% PBST-milk 4°C), then blotted with 100 ng/mL NbSym2-rabbit-Fc, recognizing the C-tag, previously conjugated to horseradish peroxidase (LighteningLink™ Novus Biologicals) according to the manufacturer’s instructions.

### Blue native polyacrylamide gel electrophoresis

RBD-multimer proteins and particles were analyzed by blue native polyacrylamide gel electrophoresis (BN-PAGE). The proteins were mixed with sample buffer and G250 loading dye and added to a 3-12% Bis-Tris NativePAGE™ gel (Life Technologies). BN-PAGE gels were run for 2 hours at 150 V using the NativePAGE™ running buffer (Life Technologies) according to the manufacturer’s instructions.

### Native Western Blotting

Expi293 supernatant samples were centrifuged and filtered (0.45-μm). Clarified supernatants were loaded and run by BN-PAGE. Gels were then transferred to 0.2-μm PVDF membrane using Xcell Blot Module under constant voltage of 25V for 1 hour. The membrane was wash and blocked (5% TBST-milk 4°C), then blotted with 50 ng/mL ACE2-Fc previously conjugated to horseradish peroxidase (LighteningLink™ Novus Biologicals) according to the manufacturer’s instructions.

## Author contributions

B.D.Q. conceived of and developed gRBD and with M.F. designed the study. B.D.Q., W.H, H.M., L.Z., and Y.G. contributed equally to the execution of these studies, with important assistance from J.C., S.P., A.O., and R.T. M.S.P., G.L.,W.L.,G.Z, and H.C. provided critical expertise and in some cases critical reagents. The manuscript was written by M.F. and B.D.Q. All authors have read the manuscript, provided comments, and approved its contents.

## Data and material availability

All requests for data or materials should be made to B.D.Q. or M.F.

## SUPPLEMENTARY MATERIALS

**Fig. S1.**
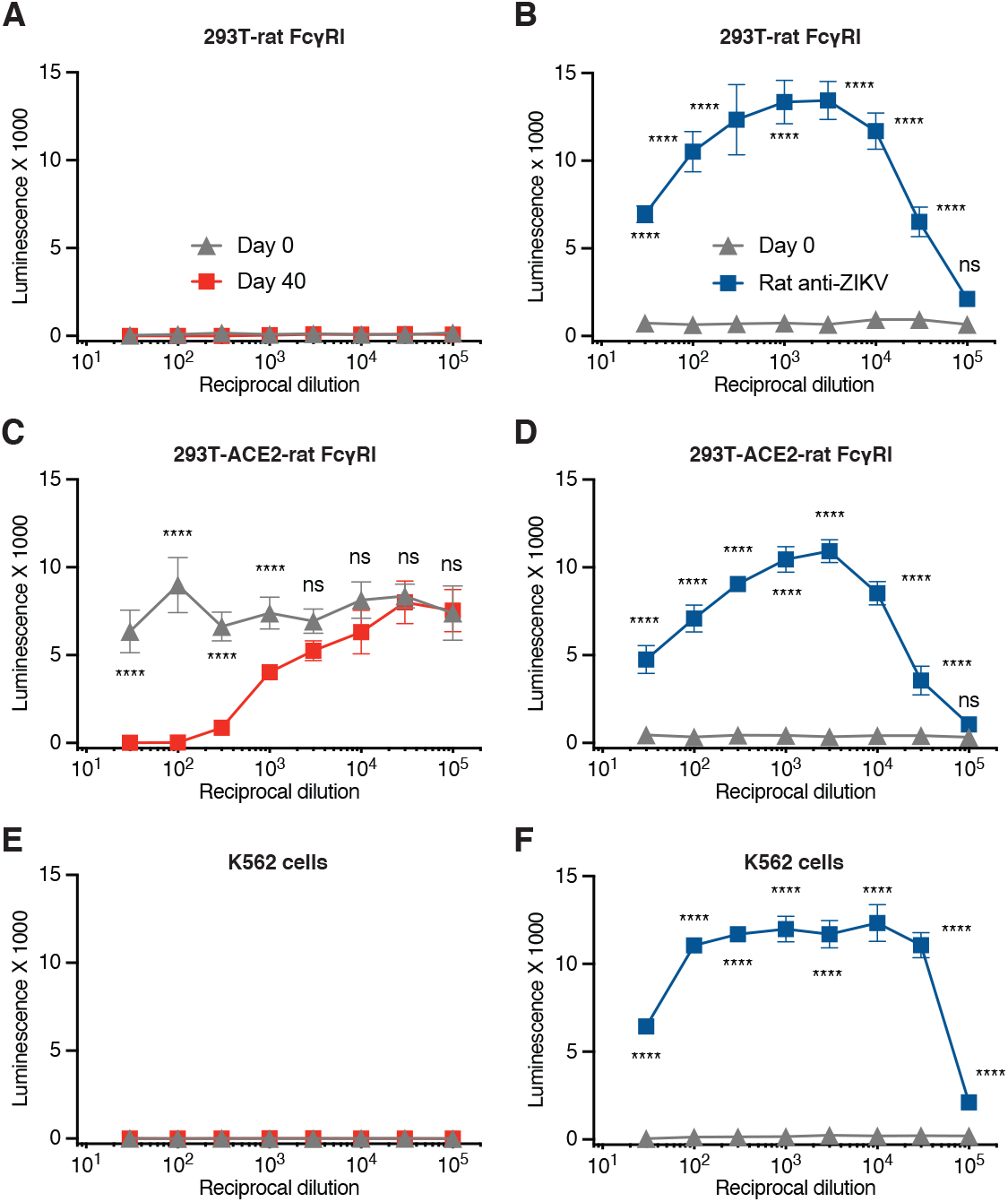
Anti-RBD antisera does not mediate antibody-dependent enhancement of SARSCoV-2 S protein-mediated entry. (**A**) HEK293T cells were transfected to express rat FcγRI and infected with SARS2-PV in the presence of the indicated dilutions of preimmune day 0 or immune day 40 antisera. (**B**) Experiments similar to those in panel A except that ZIKV VLP and anti-ZIKV antisera, also elicited in Sprague Dawley rats, were compared to preimmune sera. (**C**, **D**) Experiments identical to those in panel A and B respectively except that HEK239T cells stably expressing human ACE2 were again transfected to express rat FcγRI before incubation with SARS2-PV (**C**) or ZIKV-VLP (**D**). (**E**, **F**) Experiments identical to A and B, respectively, except that the pro-monocytic K562 cell line was incubated with SARS2-PV (E) or ZIKV-VLP (F). Error bars in panels A-F indicate the range of two experiments run in parallel. Significant differences with day-0 preimmune sera are indicated (**** indicates P<0.001 by two-way ANOVA; ns indicates not significant).

**Fig. S2.**
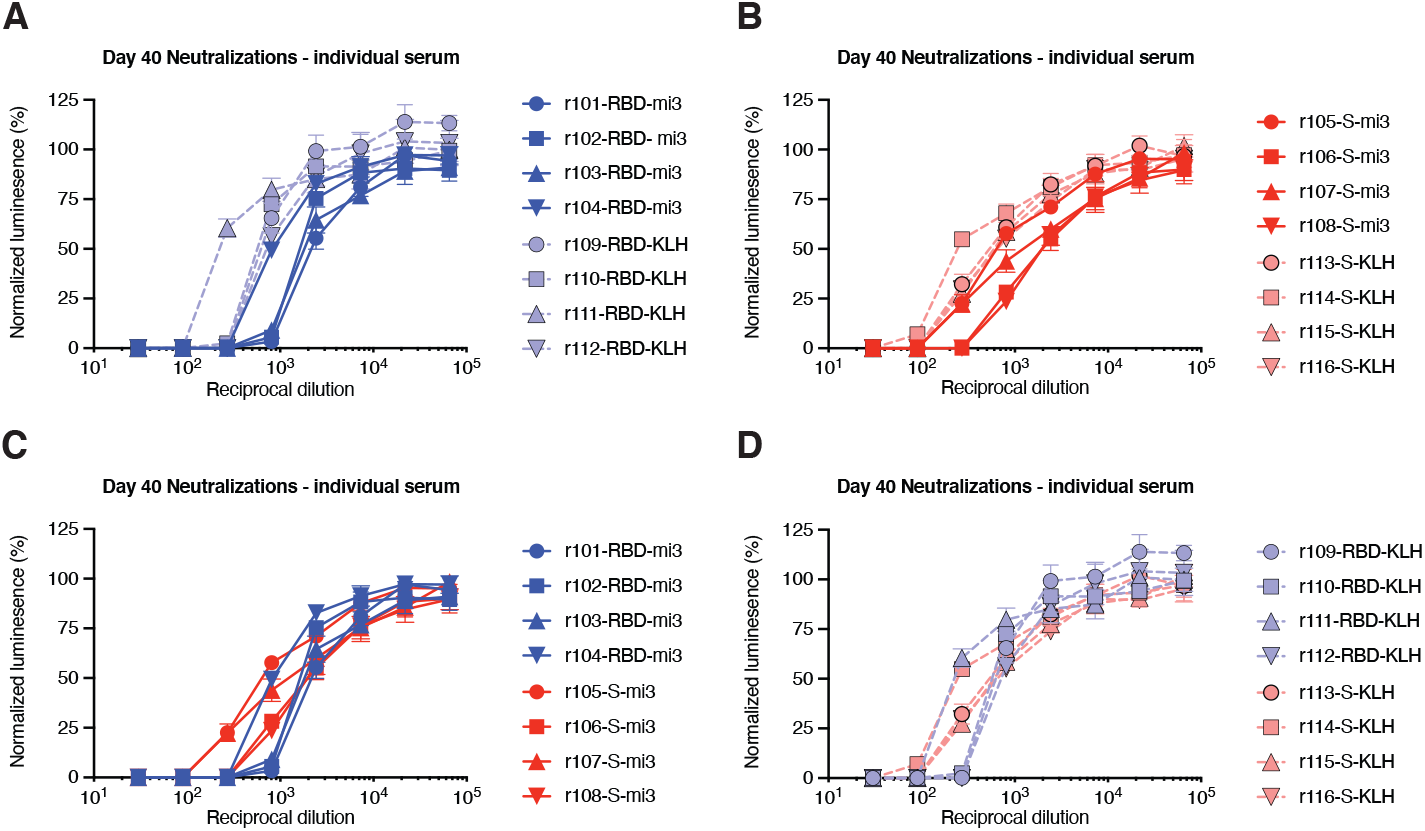
RBD/Spike mi3/KLH neutralizations for individual rats. Four female Sprague Dawley rats for each group were inoculated with either RBD-Spytag or S-protein-Spytag conguated to either Spycatcher-mi3 particles by isopeptide bond formation, or KLH by EDC. Sera were harvested at day 45 and each serum was compared for its ability to neutralize S-protein-pseudotyped retroviruses (SARS2-PV), by measuring the activity of a firefly-luciferease reporter expressed by these pseudoviruses. (**A**) RBD conjugated to mi3 vs. KLH. (**B**) S-protein conjugated to mi3 vs. KLH. (**C**) mi3 conjugates of RBD vs S-protein. (**D**) KLH conjugates of RBD vs S-protein.

**Fig. S3.**
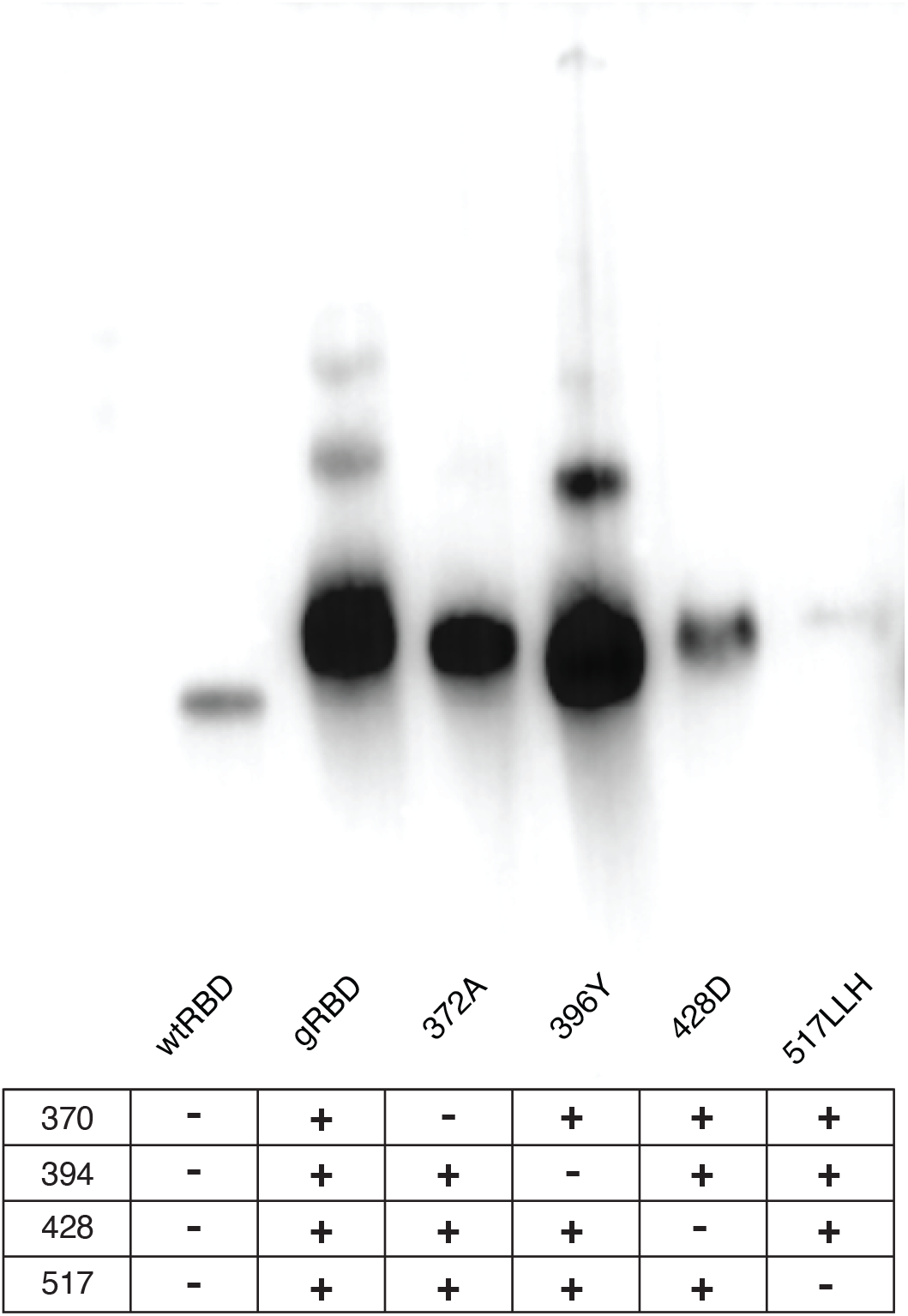
Individual glycosylation motifs promote expression of a multimeric RBD fusion protein. SARS-CoV-2 RBD variants with different combinations of glycosylation motifs, as indicated, were expressed as fusions to the C-terminus of *H. pylori* NAP 12-mer. Native western blots probed with ACE2-Fc-HRP were performed on Expi293 supernatants harvested 5 days post transfection.

**Fig. S4.**
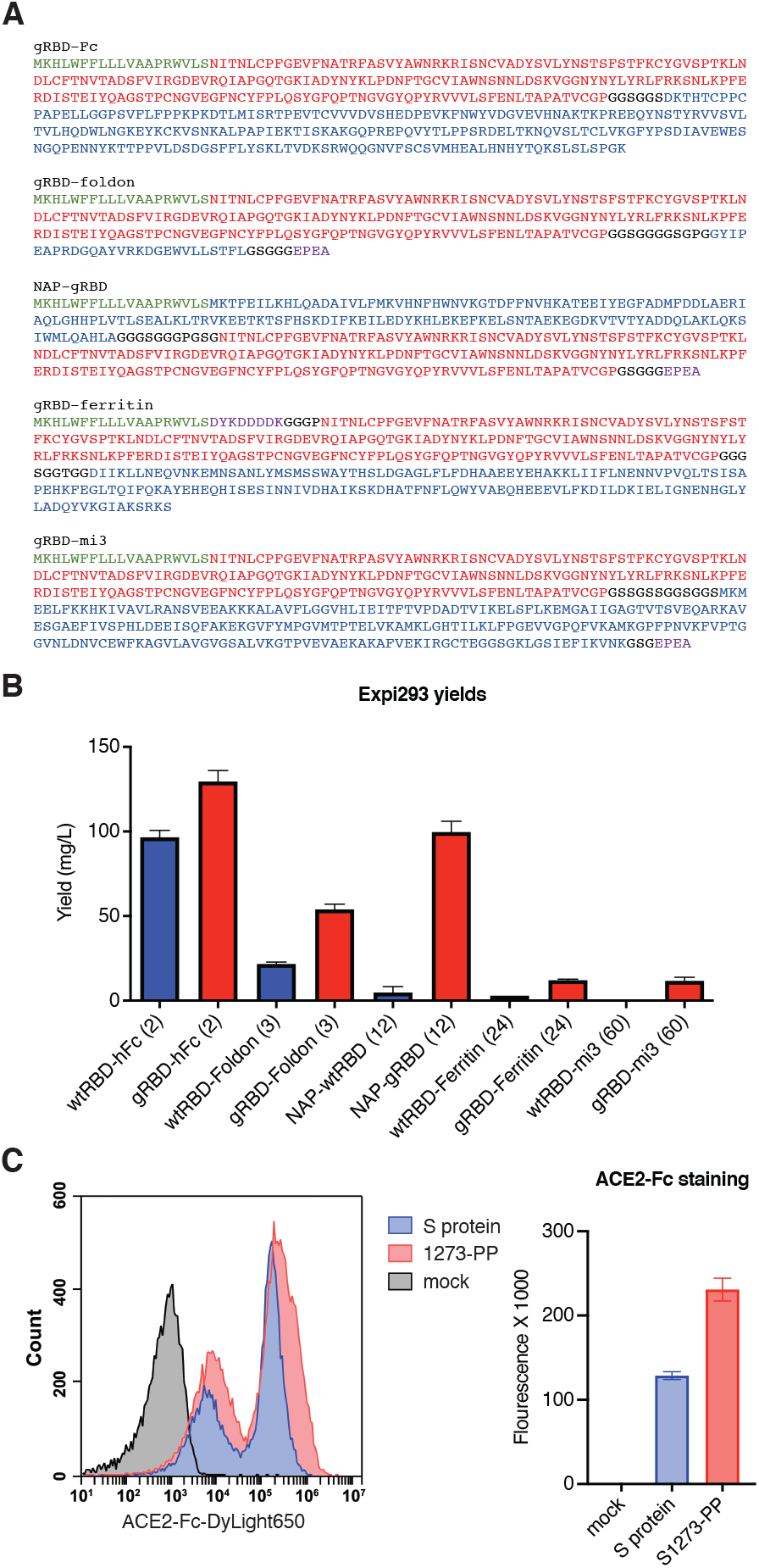
Multivalent gRBD fusion proteins express more efficiently than their wtRBD counterparts. (**A**) Complete sequences of the multivalent gRBD fusion constructs used in these studies. Green indicated signal peptide; black, linker residues; blue, multivalent carrier protein; red, gRBD; purple, affinity tag. (**B**) Yields of purified wtRBD and gRBD multimers expressed from the CMV/R vector in Expi293 cells. Values reflect a minimum of two independent transfections. Error bars indicate s.d. (**C**) Flow cytometery of 293T cells transiently transfected with SARS-CoV2 S protein in the pCAGGS vector, or S1273-PP in the CMVR vector. Expression was measured by by staining with ACE2-Fc-DyLight650. Right panel indicates mean fluorescence intensity as determine from the histogram shown in the left panel.

## REFERENCES

1. J. Cui, F. Li, Z. L. Shi, Origin and evolution of pathogenic coronaviruses. Nat Rev Microbiol 17, 181–192 (2019).

2. R. Lu, X. Zhao, J. Li, P. Niu, B. Yang, H. Wu, W. Wang, H. Song, B. Huang, N. Zhu, Y. Bi, X. Ma, F. Zhan, L. Wang, T. Hu, H. Zhou, Z. Hu, W. Zhou, L. Zhao, J. Chen, Y. Meng, J. Wang, Y. Lin, J. Yuan, Z. Xie, J. Ma, W. J. Liu, D. Wang, W. Xu, E. C. Holmes, G. F. Gao, G. Wu, W. Chen, W. Shi, W. Tan, Genomic characterisation and epidemiology of 2019 novel coronavirus: implications for virus origins and receptor binding. Lancet, (2020).

3. P. Zhou, X. L. Yang, X. G. Wang, B. Hu, L. Zhang, W. Zhang, H. R. Si, Y. Zhu, B. Li, C. L. Huang, H. D. Chen, J. Chen, Y. Luo, H. Guo, R. D. Jiang, M. Q. Liu, Y. Chen, X. R. Shen, X. Wang, X. S. Zheng, K. Zhao, Q. J. Chen, F. Deng, L. L. Liu, B. Yan, F. X. Zhan, Y. Y. Wang, G. F. Xiao, Z. L. Shi, A pneumonia outbreak associated with a new coronavirus of probable bat origin. Nature 579, 270–273 (2020).

4. V. D. Menachery, B. L. Yount, Jr., K. Debbink, S. Agnihothram, L. E. Gralinski, J. A. Plante, R. L. Graham, T. Scobey, X. Y. Ge, E. F. Donaldson, S. H. Randell, A. Lanzavecchia, W. A. Marasco, Z. L. Shi, R. S. Baric, A SARS-like cluster of circulating bat coronaviruses shows potential for human emergence. Nature medicine 21, 1508–1513 (2015).

5. N. Chen, M. Zhou, X. Dong, J. Qu, F. Gong, Y. Han, Y. Qiu, J. Wang, Y. Liu, Y. Wei, J. Xia, T. Yu, X. Zhang, L. Zhang, Epidemiological and clinical characteristics of 99 cases of 2019 novel coronavirus pneumonia in Wuhan, China: a descriptive study. Lancet, (2020).

6. W. Li, M. J. Moore, N. Vasilieva, J. Sui, S. K. Wong, M. A. Berne, M. Somasundaran, J. Sullivan, K. Luzuriaga, T. C. Greenough, H. Choe, M. Farzan, Angiotensin-converting enzyme 2 is a functional receptor for the SARS coronavirus. Nature 426, 450–454 (2003).

7. A. C. Walls, Y. J. Park, M. A. Tortorici, A. Wall, A. T. McGuire, D. Veesler, Structure, Function, and Antigenicity of the SARS-CoV-2 Spike Glycoprotein. Cell, (2020).

8. M. Hoffmann, H. Kleine-Weber, S. Schroeder, N. Kruger, T. Herrler, S. Erichsen, T. S. Schiergens, G. Herrler, N. H. Wu, A. Nitsche, M. A. Muller, C. Drosten, S. Pohlmann, SARS-CoV-2 Cell Entry Depends on ACE2 and TMPRSS2 and Is Blocked by a Clinically Proven Protease Inhibitor. Cell, (2020).

9. F. Li, Structure, Function, and Evolution of Coronavirus Spike Proteins. Annual review of virology 3, 237–261 (2016).

10. B. Coutard, C. Valle, X. de Lamballerie, B. Canard, N. G. Seidah, E. Decroly, The spike glycoprotein of the new coronavirus 2019-nCoV contains a furin-like cleavage site absent in CoV of the same clade. Antiviral research 176, 104742 (2020).

11. I. C. Huang, B. J. Bosch, W. Li, M. Farzan, P. M. Rottier, H. Choe, SARS-CoV, but not HCoV-NL63, utilizes cathepsins to infect cells: viral entry. Advances in experimental medicine and biology 581, 335–338 (2006).

12. J. K. Millet, G. R. Whittaker, Host cell proteases: Critical determinants of coronavirus tropism and pathogenesis. Virus research 202, 120–134 (2015).

13. I. Glowacka, S. Bertram, M. A. Muller, P. Allen, E. Soilleux, S. Pfefferle, I. Steffen, T. S. Tsegaye, Y. He, K. Gnirss, D. Niemeyer, H. Schneider, C. Drosten, S. Pohlmann, Evidence that TMPRSS2 activates the severe acute respiratory syndrome coronavirus spike protein for membrane fusion and reduces viral control by the humoral immune response. Journal of virology 85, 4122–4134 (2011).

14. S. Belouzard, V. C. Chu, G. R. Whittaker, Activation of the SARS coronavirus spike protein via sequential proteolytic cleavage at two distinct sites. Proceedings of the National Academy of Sciences of the United States of America 106, 5871–5876 (2009).

15. S. K. Wong, W. Li, M. J. Moore, H. Choe, M. Farzan, A 193-amino acid fragment of the SARS coronavirus S protein efficiently binds angiotensin-converting enzyme 2. The Journal of biological chemistry 279, 3197–3201 (2004).

16. F. Li, W. Li, M. Farzan, S. C. Harrison, Structure of SARS coronavirus spike receptor-binding domain complexed with receptor. Science 309, 1864–1868 (2005).

17. D. Wrapp, N. Wang, K. S. Corbett, J. A. Goldsmith, C. L. Hsieh, O. Abiona, B. S. Graham, J. S. McLellan, Cryo-EM structure of the 2019-nCoV spike in the prefusion conformation. Science 367, 1260–1263 (2020).

18. Z. Chen, L. Zhang, C. Qin, L. Ba, C. E. Yi, F. Zhang, Q. Wei, T. He, W. Yu, J. Yu, H. Gao, X. Tu, A. Gettie, M. Farzan, K. Y. Yuen, D. D. Ho, Recombinant modified vaccinia virus Ankara expressing the spike glycoprotein of severe acute respiratory syndrome coronavirus induces protective neutralizing antibodies primarily targeting the receptor binding region. Journal of virology 79, 2678–2688 (2005).

19. L. A. Jackson, E. J. Anderson, N. G. Rouphael, P. C. Roberts, M. Makhene, R. N. Coler, P. McCullough, J. D. Chappell, M. R. Denison, L. J. Stevens, A. J. Pruijssers, A. McDermott, B. Flach, N. A. Doria-Rose, K. S. Corbett, K. M. Morabito, S. O’Dell, S. D. Schmidt, P. A. Swanson, 2nd, M. Padilla, J. R. Mascola, K. M. Neuzil, H. Bennett, W. Sun, E. Peters, M. Makowski, J. Albert, K. Cross, W. Buchanan, R. Pikaart-Tautges, J. E. Ledgerwood, B. S. Graham, J. H. Beigel, R. N. A. S. G. m, An mRNA Vaccine against SARS-CoV-2 - Preliminary Report. The New England journal of medicine 383, 1920–1931 (2020).

20. L. Piccoli, Y. J. Park, M. A. Tortorici, N. Czudnochowski, A. C. Walls, M. Beltramello, C. Silacci-Fregni, D. Pinto, L. E. Rosen, J. E. Bowen, O. J. Acton, S. Jaconi, B. Guarino, A. Minola, F. Zatta, N. Sprugasci, J. Bassi, A. Peter, A. De Marco, J. C. Nix, F. Mele, S. Jovic, B. F. Rodriguez, S. V. Gupta, F. Jin, G. Piumatti, G. Lo Presti, A. F. Pellanda, M. Biggiogero, M. Tarkowski, M. S. Pizzuto, E. Cameroni, C. Havenar-Daughton, M. Smithey, D. Hong, V. Lepori, E. Albanese, A. Ceschi, E. Bernasconi, L. Elzi, P. Ferrari, Garzoni, A. Riva, G. Snell, F. Sallusto, K. Fink, H. W. Virgin, A. Lanzavecchia, D. Corti, D. Veesler, Mapping Neutralizing and Immunodominant Sites on the SARS-CoV-2 Spike Receptor-Binding Domain by Structure-Guided High-Resolution Serology. Cell 183, 1024–1042 e1021 (2020).

21. L. Liu, P. Wang, M. S. Nair, J. Yu, M. Rapp, Q. Wang, Y. Luo, J. F. Chan, V. Sahi, A. Figueroa, X. V. Guo, G. Cerutti, J. Bimela, J. Gorman, T. Zhou, Z. Chen, K. Y. Yuen, P. Kwong, J. G. Sodroski, M. T. Yin, Z. Sheng, Y. Huang, L. Shapiro, D. D. Ho, Potent neutralizing antibodies against multiple epitopes on SARS-CoV-2 spike. Nature 584, 450–456 (2020).

22. T. F. Rogers, F. Zhao, D. Huang, N. Beutler, A. Burns, W. T. He, O. Limbo, C. Smith, G. Song, J. Woehl, L. Yang, R. K. Abbott, S. Callaghan, E. Garcia, J. Hurtado, M. Parren, L. Peng, S. Ramirez, J. Ricketts, M. J. Ricciardi, S. A. Rawlings, N. C. Wu, M. Yuan, D. M. Smith, D. Nemazee, J. R. Teijaro, J. E. Voss, I. A. Wilson, R. Andrabi, B. Briney, E. Landais, D. Sok, J. G. Jardine, D. R. Burton, Isolation of potent SARS-CoV-2 neutralizing antibodies and protection from disease in a small animal model. Science 369, 956–963 (2020).

23. A. Baum, D. Ajithdoss, R. Copin, A. Zhou, K. Lanza, N. Negron, M. Ni, Y. Wei, K. Mohammadi, B. Musser, G. S. Atwal, A. Oyejide, Y. Goez-Gazi, J. Dutton, E. Clemmons, H. M. Staples, C. Bartley, B. Klaffke, K. Alfson, M. Gazi, O. Gonzalez, E. Dick, Jr., R. Carrion, Jr., L. Pessaint, M. Porto, A. Cook, R. Brown, V. Ali, J. Greenhouse, T. Taylor, H. Andersen, M. G. Lewis, N. Stahl, A. J. Murphy, G. D. Yancopoulos, C. A. Kyratsous, REGN-COV2 antibodies prevent and treat SARS-CoV-2 infection in rhesus macaques and hamsters. Science, (2020).

24. P. Chen, A. Nirula, B. Heller, R. L. Gottlieb, J. Boscia, J. Morris, G. Huhn, J. Cardona, B. Mocherla, V. Stosor, I. Shawa, A. C. Adams, J. Van Naarden, K. L. Custer, L. Shen, M. Durante, G. Oakley, A. E. Schade, J. Sabo, D. R. Patel, P. Klekotka, D. M. Skovronsky, B.-. Investigators, SARS-CoV-2 Neutralizing Antibody LY-CoV555 in Outpatients with Covid-19. The New England journal of medicine, (2020).

25. Y. He, J. Li, W. Li, S. Lustigman, M. Farzan, S. Jiang, Cross-neutralization of human and palm civet severe acute respiratory syndrome coronaviruses by antibodies targeting the receptor-binding domain of spike protein. Journal of immunology 176, 6085–6092 (2006).

26. Y. He, Y. Zhou, S. Liu, Z. Kou, W. Li, M. Farzan, S. Jiang, Receptor-binding domain of SARS-CoV spike protein induces highly potent neutralizing antibodies: implication for developing subunit vaccine. Biochemical and biophysical research communications 324, 773–781 (2004).

27. B. S. Shim, Y. C. Kwon, M. J. Ricciardi, M. Stone, Y. Otsuka, F. Berri, J. M. Kwal, D. M. Magnani, C. B. Jackson, A. S. Richard, P. Norris, M. Busch, C. L. Curry, M. Farzan, D. Watkins, H. Choe, Zika Virus-Immune Plasmas from Symptomatic and Asymptomatic Individuals Enhance Zika Pathogenesis in Adult and Pregnant Mice. mBio 10, (2019).

28. M. S. Chiofalo, G. Teti, J. M. Goust, R. Trifiletti, M. F. La Via, Subclass specificity of the Fc receptor for human IgG on K562. Cellular immunology 114, 272–281 (1988).

29. T. U. J. Bruun, A. C. Andersson, S. J. Draper, M. Howarth, Engineering a Rugged Nanoscaffold To Enhance Plug-and-Display Vaccination. ACS nano 12, 8855–8866 (2018).

30. Y. Hsia, J. B. Bale, S. Gonen, D. Shi, W. Sheffler, K. K. Fong, U. Nattermann, C. Xu, P. S. Huang, R. Ravichandran, S. Yi, T. N. Davis, T. Gonen, N. P. King, D. Baker, Design of a hyperstable 60-subunit protein dodecahedron. [corrected]. Nature 535, 136–139 (2016).

31. H. Duan, X. Chen, J. C. Boyington, C. Cheng, Y. Zhang, A. J. Jafari, T. Stephens, Y. Tsybovsky, O. Kalyuzhniy, P. Zhao, S. Menis, M. C. Nason, E. Normandin, M. Mukhamedova, B. J. DeKosky, L. Wells, W. R. Schief, M. Tian, F. W. Alt, P. D. Kwong, J. R. Mascola, Glycan Masking Focuses Immune Responses to the HIV-1 CD4-Binding Site and Enhances Elicitation of VRC01-Class Precursor Antibodies. Immunity 49, 301–311 e305 (2018).

32. D. W. Kulp, J. M. Steichen, M. Pauthner, X. Hu, T. Schiffner, A. Liguori, C. A. Cottrell, C. Havenar-Daughton, G. Ozorowski, E. Georgeson, O. Kalyuzhniy, J. R. Willis, M. Kubitz, Y. Adachi, S. M. Reiss, M. Shin, N. de Val, A. B. Ward, S. Crotty, D. R. Burton, W. R. Schief, Structure-based design of native-like HIV-1 envelope trimers to silence non-neutralizing epitopes and eliminate CD4 binding. Nature communications 8, 1655 (2017).

33. S. Meier, S. Guthe, T. Kiefhaber, S. Grzesiek, Foldon, the natural trimerization domain of T4 fibritin, dissociates into a monomeric A-state form containing a stable beta-hairpin: atomic details of trimer dissociation and local beta-hairpin stability from residual dipolar couplings. Journal of molecular biology 344, 1051–1069 (2004).

34. G. Zanotti, E. Papinutto, W. Dundon, R. Battistutta, M. Seveso, G. Giudice, R. Rappuoli, C. Montecucco, Structure of the neutrophil-activating protein from Helicobacter pylori. Journal of molecular biology 323, 125–130 (2002).

35. M. Kanekiyo, C. J. Wei, H. M. Yassine, P. M. McTamney, J. C. Boyington, J. R. Whittle, S. S. Rao, W. P. Kong, L. Wang, G. J. Nabel, Self-assembling influenza nanoparticle vaccines elicit broadly neutralizing H1N1 antibodies. Nature 499, 102–106 (2013).

36. L. E. Gralinski, R. S. Baric, Molecular pathology of emerging coronavirus infections. The Journal of pathology 235, 185–195 (2015).

37. S. T. Reddy, A. J. van der Vlies, E. Simeoni, V. Angeli, G. J. Randolph, C. P. O’Neil, L. K. Lee, M. A. Swartz, J. A. Hubbell, Exploiting lymphatic transport and complement activation in nanoparticle vaccines. Nature biotechnology 25, 1159–1164 (2007).

38. M. O. Oyewumi, A. Kumar, Z. Cui, Nano-microparticles as immune adjuvants: correlating particle sizes and the resultant immune responses. Expert review of vaccines 9, 1095–1107 (2010).

39. L. Zhang, W. Ji, S. Lyu, L. Qiao, G. Luo, Tet-Inducible Production of Infectious Zika Virus from the Full-Length cDNA Clones of African- and Asian-Lineage Strains. Viruses 10, (2018).

